# The impacts of light and location on reproductive timing in the Orbicella species complex

**DOI:** 10.64898/2026.09.14.751534

**Authors:** Elizabeth A. Boville, Leanne M. Ulmer, Nancy Knowlton, Don Levitan, Oren Levy, Alina Szmant, Mónica Medina

## Abstract

Coral spawning events rely on tight synchronicity of gamete release to increase fertilization success by increasing gamete density in the water. Differences in spawning time can cause temporal reproductive isolation and minimize hybridization with closely related coral species. Sexual reproductive success is important as it generates the genetic diversity necessary for corals to survive large scale environmental stressors such as heat waves and bleaching events. The coordination of coral spawning events within a population maintains species boundaries and increases ecosystem-wide genetic diversity. Here we generate a database of spawning events in conjunction with depth and geographic location of the three species in the *Orbicella* species complex to assess variability in reproductive timing. The dataset, gathered from published papers, restoration practitioners, scientist throughout the Caribbean, contains almost three decades of data and includes 265 reproductive events. Our findings indicate that some aspects of each spawning season (month and day after the full moon) are influenced by latitude whereas others, such as spawning time in relation to local sunset, may exhibit variation based on geographic location. Combined, our results suggest that the reproductive timeframe of *Orbicella* spp. is triggered by factors associated with latitudinal gradients, such as the amount of energy from the sun, but the specific time of gamete release may be connected to other environmental factors that occur at a local scale or vary year-to-year.

## Introduction

The importance of coral reefs cannot be understated. These rainforests of the ocean provide vital habitat for fish and other marine taxa, protect shorelines from waves and storm damage, and create sources of income for local economies through fishing and tourism (Costanza, 1997; Carpenter et al., 2008; Harborne et al, 2017). However, the threats these fragile ecosystems face are monumental on a global scale (Carpenter et al., 2008; Harborne et al, 2017; Hughes et al., 2018; Andrello et al., 2022). Mass bleaching events (Hoegh-Guldberg 1999; Donner et al., 2005; Baker et al., 2008, Hughes et al., 2017; Lenz et al., 2025) often due to increased seawater temperature (Hoegh-Guldberg, 1999; Bruno et al., 2007), higher storm intensity (Hughes, 1994), overfishing (Huges, 1994), and disease outbreaks (Bruno et al., 2007) threaten coral health worldwide and greatly reduce reef size and biodiversity (Bellwood et al., 2004; Carpenter et al., 2008; Andrello et al., 2022).

Conservation efforts to preserve and repopulate reefs are in motion, focusing primarily on coral fragmentation and rearing coral larvae to outplant into their natural environments (Baums et al., 2019). These efforts are productive but challenging as many scleractinian corals are slow-growing and are hard to raise *ex-situ* (Baums et al., 2019). Clonal colonies produced through fragmentation can be more severely impacted by bleaching and disease outbreaks due to reduced genetic diversity as they are a product of asexual reproduction, limiting the efficacy of this restoration practice (Baums et al., 2019; Chan, Peplow & van Oppen, 2019). Raising larvae produced through sexual reproduction can provide more resilient populations due to increased genetic variation as different genotypes can cope with diverse environmental conditions (Harborne et al, 2017; Baums et al., 2019; Thomas et al., 2024; Lenz et al., 2025). Many coral species reproduce in annual broadcast spawning events in which most colonies in a location release gamete bundle containing eggs and sperm into the water column at a precise date and time (Babcock et al., 1986; Szmant, 1986; Harrison, 2011; Levitan et al., 2011). Levitan et al. (2011) showed that there is also a genetic component to spawning times as individual colonies displayed minimal variation between years. Studies indicate that the tight synchronization and high gamete density increases fertilization success while reducing inter-species interactions (Babcock et al., 1986; Levitan et al., 2004; Lacharmoise & Köksal, 2006; Van Woesik, 2006; Kaniewska et al., 2015).

The precise date of spawning events in many Caribbean scleractinian coral species is tied to lunar cycles, with most species spawning in the week after the full moon between July and September (Fadlallah, 1983; Szmant, 1991; Vize et al., 2005; Monfared et al., 2023). Gamete release time during those months varies between species but is thought to be triggered by day length as many species spawn at a precise time after sunset (Szmant, 1991; Levitan et al., 2004; Vize et al., 2005; Levitan et al., 2011; Monfared et al., 2023). Such a small spawning window makes it challenging to collect gametes and monitor spawning behavior between years. Regional variation in spawning timing has been noted based on latitude as well as sea surface temperature shifts, wind speed and light availability (Kaniewska et al., 2015; Sakai et al., 2020; Monfared et al., 2023). Spawning desynchronization has been observed due to artificial light at night (ALAN), a measurement of anthropogenic light pollution (Ayalon et al., 2021; Davies et al., 2023), and increased seawater temperature during the reproductive cycle (Shlesinger & Loya, 2019; Furukawa et al., 2020; Levy et al., 2020; Tamir et al., 2020; Fobert, 2023). Changes to the date and time of gamete release may be significant enough to form reproductive barriers by creating genetically isolated populations and reducing fertilization success rates (Levitan et al., 2004; Furukawa et al., 2020; Levy et al., 2020; Ayalon et al., 2021; Davies et al., 2023, Fobert, 2023).

Here we focus on *Orbicella annularis, O. faveolata*, and *O. franksi*, three broadcast-spawning species comprising the *Orbicella* species complex. These endangered corals are prominent reef-builders throughout the Caribbean (Budd & Klaus, 2001; Budd, 2010; Prada et al., 2016). *Orbicella* spp. are known to inhabit light-specific depth ranges, with *O. annularis* typically found at shallow depths (< 20m), *O. franksi* found at deeper depths (10-50m), and *O. faveolata* found at a variety of depths spanning 2-30m (Knowlton et al., 1992; Fukami et al., 2004; Prada et al., 2016; Egan et al., 2021; Prada et al., 2024). These depth ranges can vary slightly, as colonies at different depths and water turbidities exhibit distinct morphologies in response to light availability and water quality (Knowlton et al., 1992; Fukami et al., 2004; López-Victoria et al., 2015; Prada et al., 2016; Pizarro et al., 2017; Egan et al., 2021; López-Londoño et al., 2021). *Orbicella* spp. are therefore highly prevalent throughout the Caribbean and comprise a key study system to assess reproductive timing at a variety of geographic locations and depths. A widespread comparison of spawning behavior across the three *Orbicella* species would allow for insights as to what habitats are most conducive to stable reproductive timing, and how habitat and environmental change may influence spawning behavior.

*Orbicella* reproduce in synchronous mass broadcast-spawning events in which gamete bundles containing eggs and sperm are released into the water column at a precise time and date (Knowlton et al., 1997; Vize et al., 2005; Levitan et al., 2004; Levitan et al., 2011). Their reproductive cycle typically begins in May with increased water temperature triggering gametogenesis, particularly in northern latitudes (Szmant, 1991; Van Veghel, 1994; Sánchez et al., 1999; Van Woesik et al., 2006). Spawning occurs in August or September, with occasional events in July or October (Szmant, 1991; Knowlton et al., 1997; Szmant et al., 1997; Sánchez et al., 1999). The three species exhibit differences in their spawning time with *O. franksi* spawning 1-2 hours earlier than the other two species although the time of gamete release has been shown to vary between geographic locations (Knowlton et al., 1997; Sánchez et al., 1999; Levitan et al., 2004; Levitan et al., 2011; Fogarty et al., 2012). The observed differences in spawning time are thought to create species boundaries as there is some evidence of *ex-situ* hybridization between*O. annularis* and *O. franksi* (Levitan et al., 2004; Levitan et al., 2011). The depth range of these two species, combined with their observed difference in spawning time, is thought to reduce the possibility of hybridization *in-situ* (Levitan et al., 2004; Levitan et al., 2011; Fogarty et al., 2012).

While general trends in *Orbicella* spawning behavior have been recorded, it is not well understood how spawning time may differ between sites and local habitats<u>. M</u>uch of the data stored on online databases <u>have not examined site-specific temporal trends of increasing</u> temperature and light pollution. This study aims to utilize spawning time observations for *Orbicella* spp. to identify variation in reproductive behavior to examine associations between spatiotemporal factors and variation in spawning time. Our curated dataset spans over three decades of spawning information over the entire biogeographical range of the three *Orbicella* species. Using this dataset, we assessed variation in peak spawning time relative to sunset, days after the full moon (DAFM), and spawning month, and found that latitudinal gradients and geographic location can influence spawning behavior. Unraveling these trends in reproductive timing in *Orbicella* spp. can lead to more effective restoration practices and help us understand factors that may impact reproductive success. A widespread comparison of the spawning behavior across the three species will allow us to understand the conditions that are conducive to stable reproductive timing, and how habitat and environmental change may influence spawning behavior.

## Methods

### Data acquisition

We combined data from published literature and unpublished records primarily from researchers, Non-Governmental Organizations (NGOs), and coral restoration groups into a comprehensive *Orbicella* spp. spawning dataset (Supplement Table 1). Observations were recorded as calendar day, geographic location, and spawning time. Additional information such as depth, weather conditions, and spawning amount was noted when available. Sunrise, sunset, moonrise, moonset, and moon peak data were calculated using the US Navy Astronomical Applications Department’s Complete Sun and Moon Data for One Day database. To ensure that the values were accurate, the latitude and longitude for each location were used in addition to the local date and time. Time zones and daylight savings time were accounted for based on the geographic location. Spawning time in hours post-sunset (HPS) and days after the full moon (DAFM) were calculated based on the values found for sunset and full moon date.

### Statistical Analysis

All data analysis was done in R v4.2.2 (R Core Team, 2022). ANOVA was used to assess differences in mean spawning time between species. Generalized linear models (GLMs) were used to calculate correlations between spawning timeframe, latitude, and depth, then plotted with ggplot2. Map generation was conducted using ggplot2 and naturalearth packages to visualize the ranges of spawning timing at different geographic locations for each species. Heat maps of datapoints were plotted over the Caribbean locations (Wickham & Sievert, 2016; Massicotte, 2023) to visualize latitudinal and geographic variation between sites and significance was assessed using GLMs.

## Results

### Dataset Overview

A total of 265 *Orbicella* spawning events from 13 Caribbean countries and 73 distinct dive sites were analyzed in this study spanning over three decades (1991-2024). We identified 81 observations for *O. annularis*, 149 for *O. faveolata*, and 35 for *O. franksi*. One *O. franksi* datapoint was omitted from the analysis as the listed spawning time was substantially later than the average and corresponds with that of *O. faveolata*, suggesting potential species misidentification. The datapoint contained one colony, and species identification could not be confirmed due to lack of visual verification. Each spawning observation consists of a single spawning event and contains 1-50+ individual colonies (Supplement Table 1). Recorded locations range in latitude from approximately 28° in the Northern Caribbean (Florida, Bahamas, Flower Garden Banks) to approximately 9° in the Southern Caribbean (Colombia, Curaçao) (Supplement Table 1).

To define the typical spawning time, day, and month, we primarily used observations from mass spawning events with greater than three colonies studied and >10% of colonies released gamete bundles. These datapoints are defined as “Peak” and “Off-Peak”, with Peak spawning events indicating spawning dates with the maximum spawning during the observation period and Off-Peak indicating spawning dates where mass spawning occurred but less spawning (% of colonies) than on the Peak date. We also incorporated spawning events with a small number or % colonies spawned as “Minimal” spawning observations to thoroughly incorporate instances of *Orbicella* reproductive events.

### Characterization of typical spawning timeframe

Our analysis found that *O. annularis* and *O. faveolata* typically spawn at 3.57 and 3.43 hours post-sunset (HPS) respectively (Table 1). In *O. annularis*, we found that the average and median spawning time was comparable to that of *O. faveolata*, varying by six minutes or less (Table 1). These two species also have the same maximum recorded average spawning time, 4.13 HPS. The minimum average spawning time for *O. faveolata* is 1.25 hours earlier than that of *O. annularis* (Table 1). The typical spawning time of *O. franksi* significantly differs from that of the other two species, with an average time of gamete release at 2.19 HPS (Table 1). The range of this species is larger, spanning from 0.25-3.69 HPS (Table 1). While the datapoints for the shallow-dwelling *O. annularis* are centered around the mean, the other two species exhibit more variation in their observed time of gamete release.

Our data show that the generalist species *O. faveolata* is observed to spawn from 1.25-4.13 HPS (Table 1), overlapping with the average value for *O. franksi* as well as that of *O. annularis* (Fig. 1). Spawning amount did not indicate any significant difference in spawning time and Peak, Off-Peak, and Minimal spawning observations were used to assess overall spawning hour behavior.

**Fig. 1.**
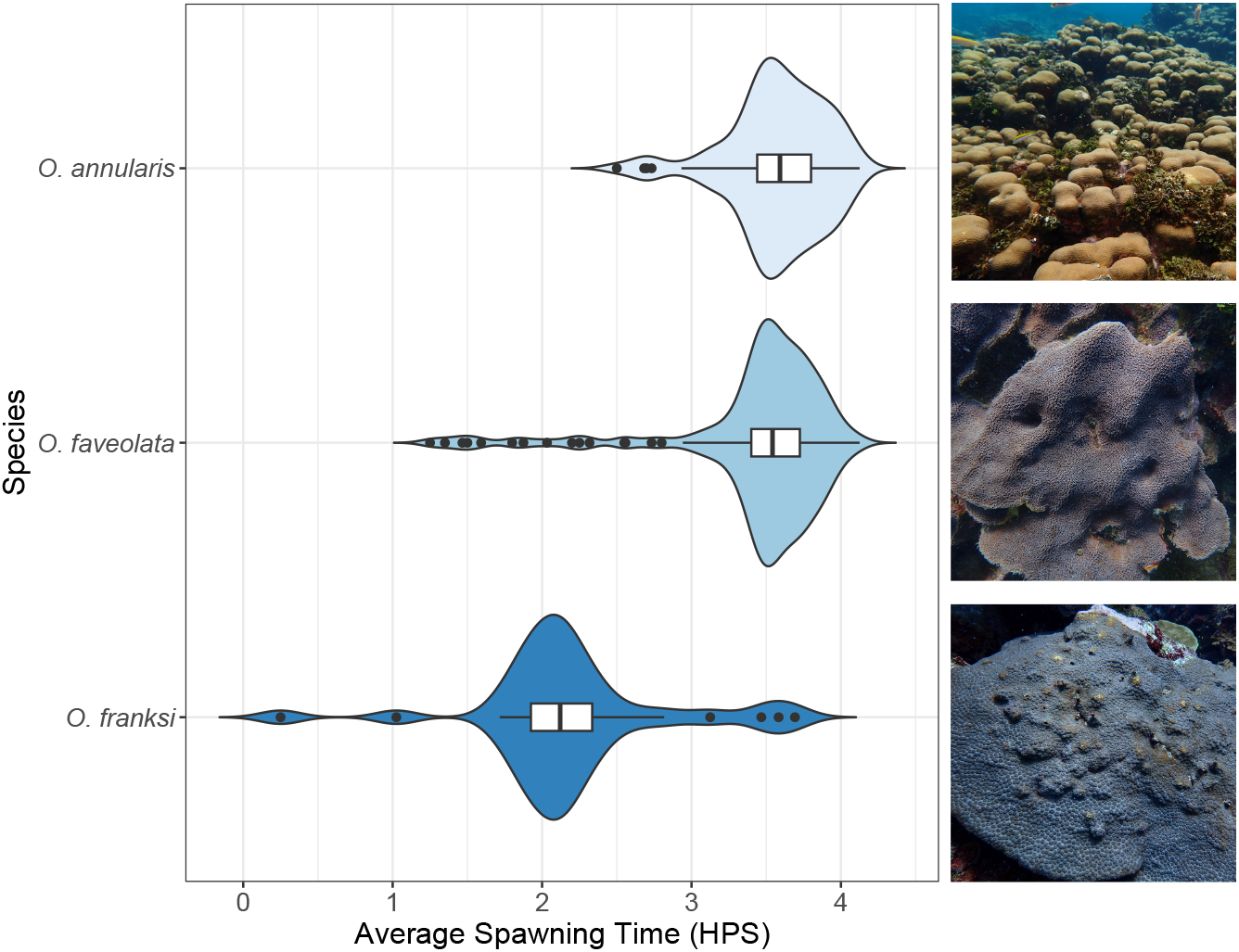
The average spawning time in hours post-sunset (HPS) for each observation recorded in our database with *Orbicella annularis* typically spawning 3.57 HPS (n=81), *O. faveolata* spawning 3.43 HPS (n=149), and *O. franksi* spawning 2.19 HPS (n=35). Values specified in Table 1.

**Table 1.** Spawning time (hours post-sunset) averaged for each observation for three *Orbicella species*. Mean values include standard deviation.

*O. annularis* n= 81, *O. faveolata* n=149, *O. franksi* n=35.
| Species | Mean | Median | Min. | Max. |
| --- | --- | --- | --- | --- |
| *O. annularis* | 3.57 +/- 0.33 | 3.59 | 2.5 | 4.13 |
| *O. faveolata* | 3.43 +/- 0.55 | 3.54 | 1.25 | 4.13 |
| *O. franksi* | 2.19 +/- 0.64 * | 2.12 | 0.25 | 3.69 |

Spawning events occur within two weeks after the full moon, with most observations for all three species were reported 6-9 DAFM (Fig. 2). *O. annularis* and *O. faveolata* typically have major (Peak) spawning events 7 DAFM while *O. franksi* has a higher frequency of Peak spawning observations 8 DAFM (Fig. 2). In *O. annularis* we identified a small number of observed Minimal spawning events as early as 4 and 5 DAFM and as late as 10 DAFM (Fig. 2a). Off-Peak and Minimal spawning records show that smaller spawning events occur leading up to and after the Peak spawning night, particularly in *O. annularis* and *O. faveolata* (Fig. 2a,b). The spawning day relative to the full moon in *O. franksi* occurs in a narrower window than the other two species, with a high frequency of observations of mass spawning events occurring 7-8 DAFM. This merits further investigation due to the fewer overall number of observations

**Fig. 2.**
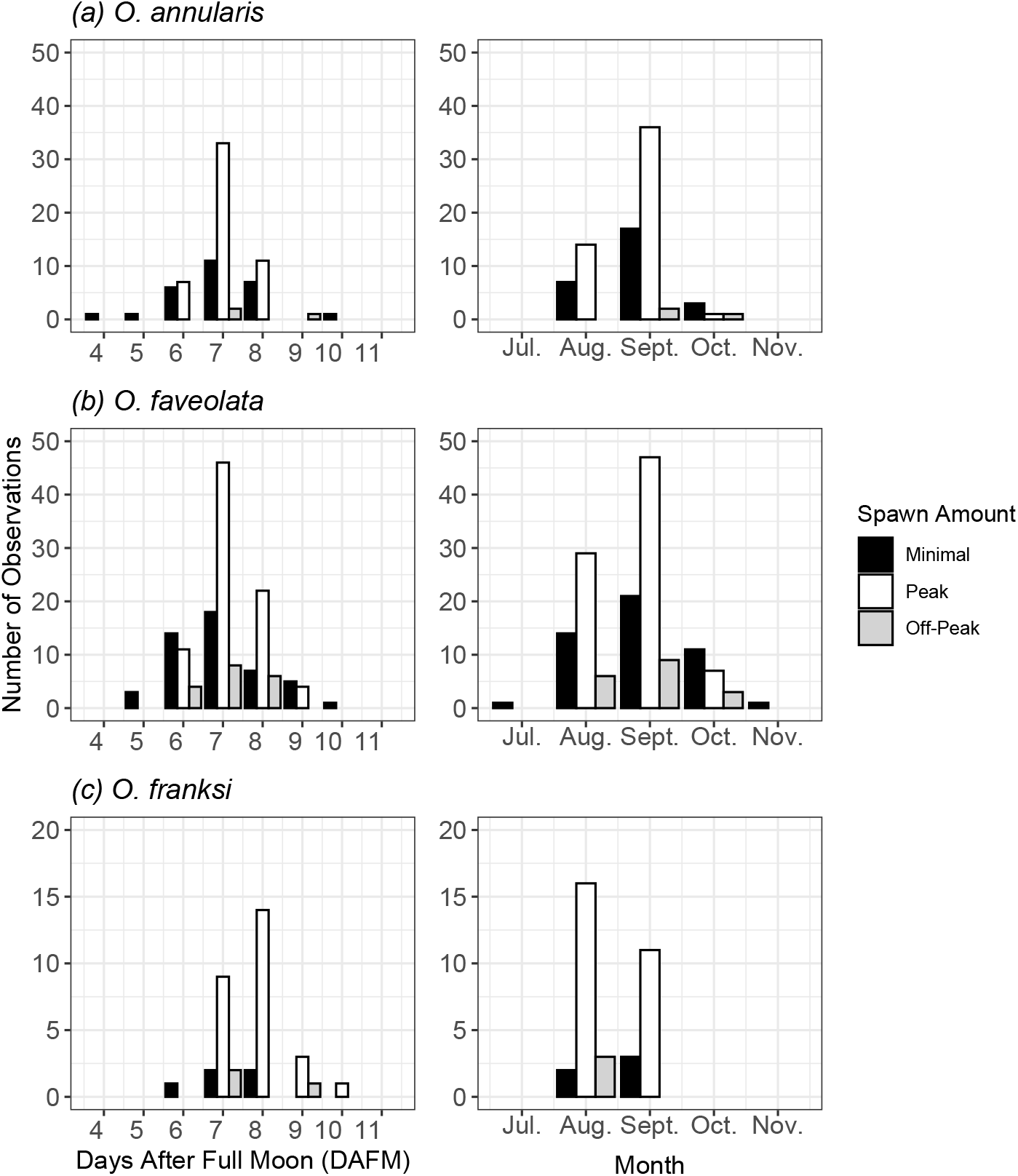
The number of distinct observations for (a) *Orbicella annularis* (n= 81), (b) *O. faveolata* (n=149) and (c) *O. franksi* (n=35) in relation to day after the full moon (left) and month (right). Each observation may contain many coral colonies (1-50+) and indicates one spawning event.

Our assessment of all recorded observations found that Peak spawning in *Orbicella* spp. typically occurs in August or September, with a small number of spawning events in July, October, and November. We found that *O. annularis* and *O. faveolata* primarily have Peak spawning dates in August and September, with 77 and 125 recorded observations respectively (Fig. 2a,b). *O. franksi* was observed to spawn primarily in August, with 25 recorded spawning events that month (Fig. 2c). There were no records of *O. annularis* spawning in July, and no records of *O. franksi* spawning in October (Fig. 2a,c). *O. faveolata* has been observed to spawn primarily in August and September but was observed spawning in all five months (Fig. 2b).

Average spawning time (HPS) displayed no significant change based on day (DAFM) or month (Fig. 3). A slight increase in average spawning time approaching the most common spawning days (7-8 DAFM) was observed in *O. annularis* and *O. faveolata*, followed by a decrease in average spawning time after the primary spawn night (Fig. 3a,b).

**Fig. 3.**
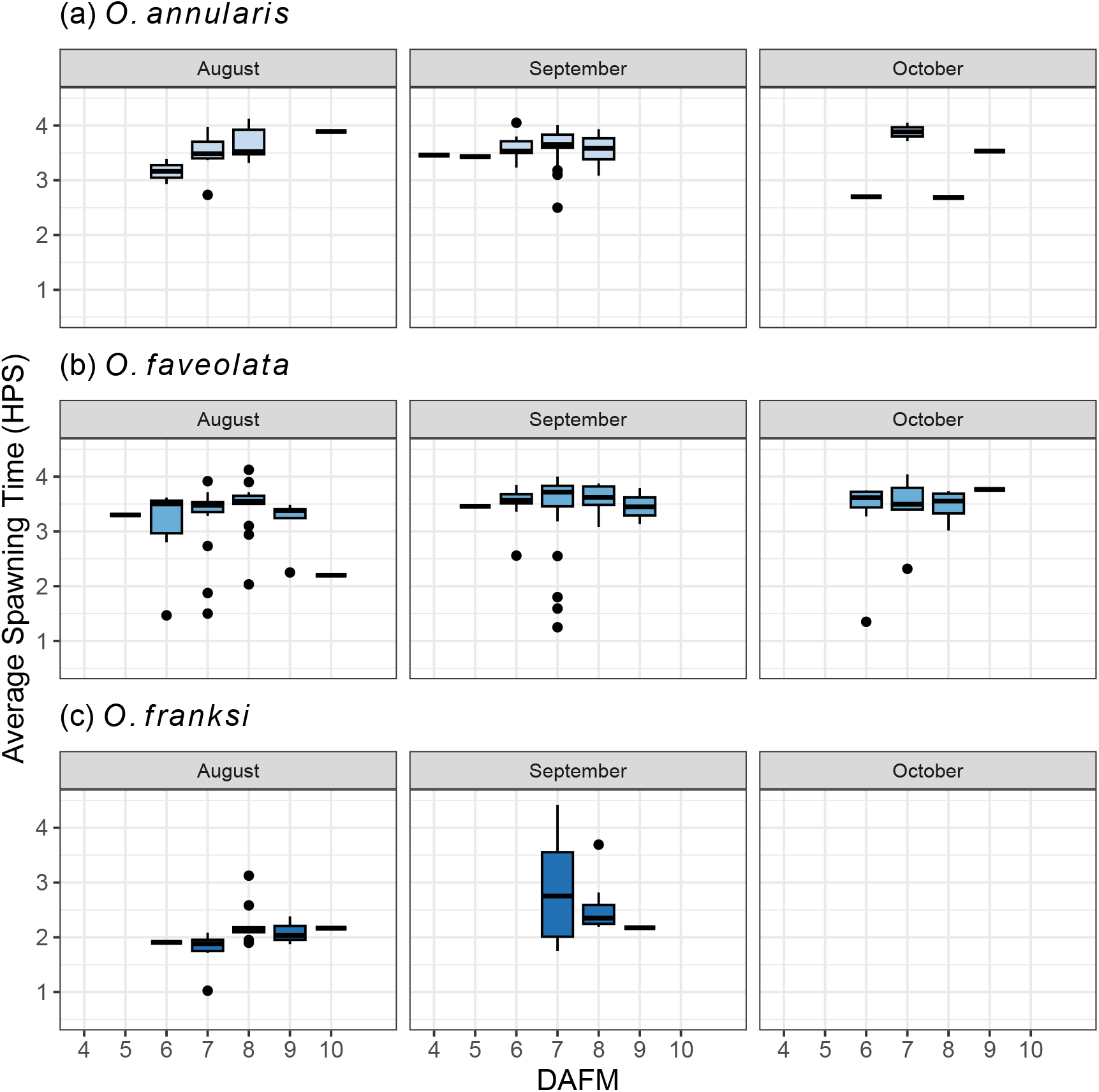
Average spawning time (HPS) of mass spawning events in (a) *Orbicella annularis* (n=54), (b) *O. faveolata* (n=101) and (c) *O. franksi* (n=30) in relation to day after the full moon (DAFM) and spawn month. Each observation may contain many coral colonies (3-50+) and indicates one spawning event.

### Geographic Location and Latitudinal Variation

Our dataset contains measurements from *Orbicella* spp. colonies found throughout the Caribbean, with a latitude range of approximately 9 to 28 decimal degrees. All three of these species were found to inhabit the full range of latitudes in our study.

To further understand the observed variation in spawning time between datapoints, we visualized each spawning observation based on location and used generalized linear models (GLMs) to assess correlation between latitude and spawning timing. Many dive sites were studied over several years, allowing for an analysis of variation between observations at each location. In all three species there was no significant correlation between latitude and spawning time (HPS) (Fig. 4a-c). In *O. annularis* and *O. faveolata* day after the full moon (DAFM) (Fig. 4d-f) was significantly correlated with latitude (p < 0.05). We found that the spawning hour (HPS) of *O. annularis* was generally consistent at all dive sites, with no location spawning significantly earlier or later than other locations (Fig. 4a). Although we collected fewer datapoints for the deeper-dwelling *O. franksi*, some regions, specifically Flower Garden Banks and Panama, tend to spawn earlier than other regions (Fig. 4c). Records from Florida and the Bahamas, locations with similar latitudes, exhibit variation in spawning time despite their close proximity for all three coral species (Fig. 4a-c). Spawn month is correlated with latitude, with corals at higher latitudes typically spawning in August and corals at lower latitudes spawning inSeptember (p < 0.01) (Fig. 5g-i). The gradient in spawn month, with lower latitudes spawning later, is present throughout the timeframe of our gathered and is most prominently observed in *O. faveolata* as some locations record spawning events in October.

**Fig. 4.**
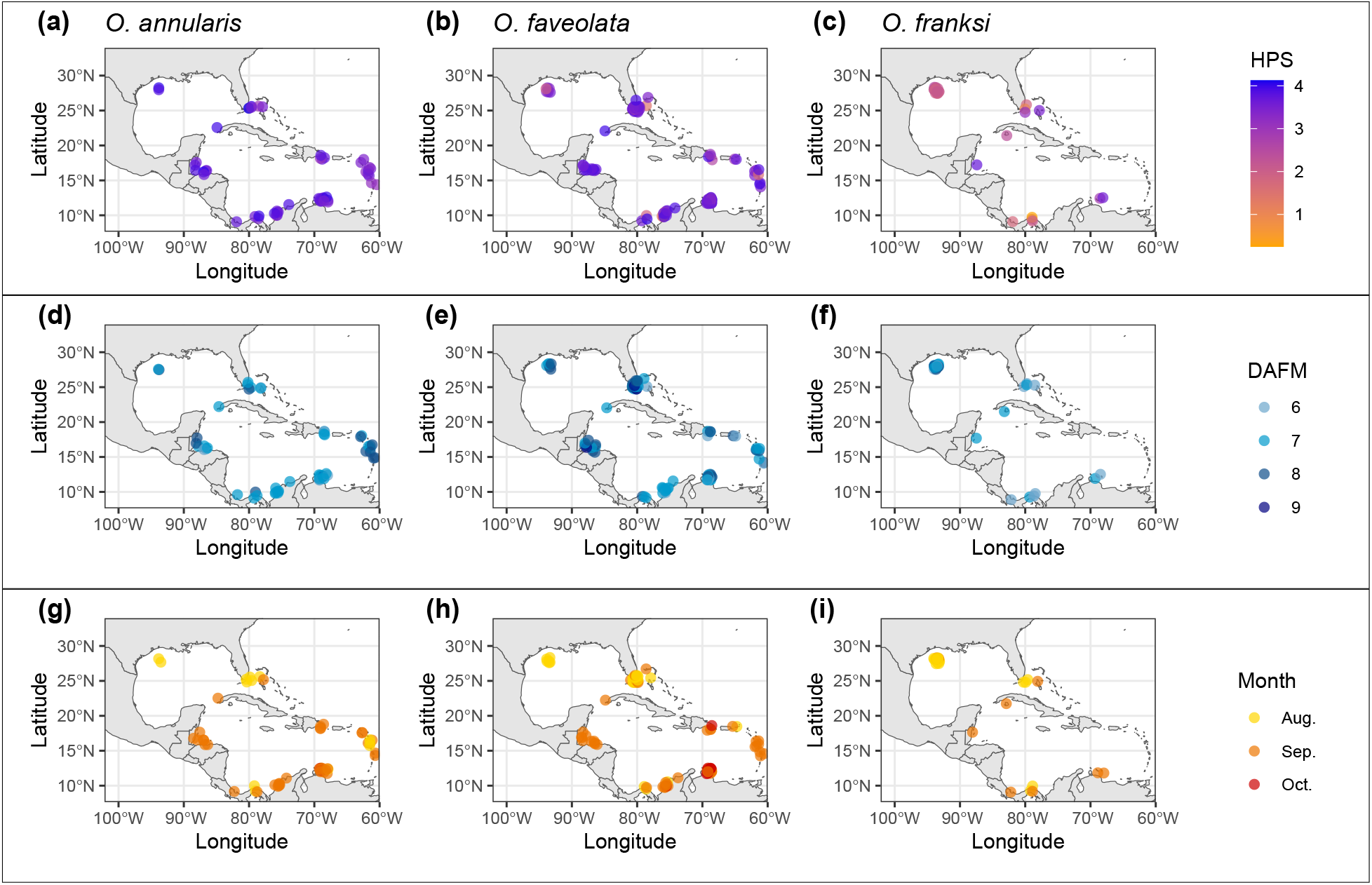
Spawning overview maps visualizing spawning time in relation to geographic location for *O. annularis* (n=54), *O. faveolata* (n=101), and *O. franksi* (n=30) from 1991-2024. Spawning time in hours post-sunset (HPS) (a-c) is not significant. Spawning day after the full moon (DAFM) (d-e) in *O. annularis* and *O. faveolata* is significantly correlated with latitude (p<0.05). Spawning month (g-i) is significant in all three species (p<0.05). Each observation may contain many coral colonies (3-50+) and indicates one spawning event. Points are jittered using R to show overlapping records and are displayed in an approximate location based on the coordinates recorded by the observers. Significance calculated using generalized linear models.

## Discussion

### Spawning Timeframe and Within-Species Variability

Here we establish the typical spawning date and time for the three extant *Orbicella* species. We found that *O. annularis* and *O. faveolata* are more likely to spawn in synchrony, with major spawning events typically occurring seven days after the full moon (DAFM) in the months of August or September. Our results indicate that *O. franksi* generally spawns 7-8 DAFM in August, however this range appears to be flexible as prior studies have shown that *O. franksi* spawning events in Panama occurred 7-8 DAFM in 1994 and 1995 and 5-6 DAFM 2002-2009 (Knowlton et al., 1997; Levitan et al., 2004). We observed more variability of spawning time in *O. faveolata* in comparison with the other two species with many observations recording earlier spawning timepoints that overlap with the mean spawn time in *O. franksi* as well as later recordings overlapping with *O. annularis*. The observed variation in spawning time within species and at specific geographic locations suggests that gamete release time is impacted by external factors that change year-to-year. Both Knowlton et al., (1997) and Levitan et al., (2004) observed that *O. franksi* spawns at a higher rate in August, aligning with our findings. *O. faveolata* exhibits more variation in spawning month than the other two species, with a small number of colonies observed spawning in July and November. The broader time (HPS), date (DAFM), and month range is possibly a result of the generalist life strategy of *O. faveolata* as this species is known to inhabit a broad range of depths and exhibits morphological differences (e.g., colony shape, color) based on location on the reef (Budd & Klaus, 2001; López-Londoño et al., 2021). This phenotypic plasticity may extend to spawning time and date, reflecting a larger sensitivity to environmental factors that are involved in reproductive timing (Van Woesik, Lachrmoise & Köksal, 2006; Gilmour, Speed & Babcock, 2016; Sakai et al., 2020; Sheridan et al., 2025).

Combined, the difference in spawning date and time, particularly between *O. annularis* and *O. franksi* are indicative of temporal reproductive isolation between these two species as they have been shown to hybridize in *ex-situ* conditions with simulated synchronous spawning events with some success (Levitan et al., 2004). The consistent maintenance of different spawning timing throughout the Caribbean reinforces the boundary between these two species. The presence of temporal reproductive barriers is further supported by variance in spawning time and date in *O. faveolata*. We identified that *O. faveolata* exhibits overlapping spawn date and time with both *O. annularis* and *O. franksi*. Levitan et al. (2004) observed no cross fertilization in *O. faveolata* with either species, suggesting that strong non-temporal reproductive barriers are present (Sánchez et al., 1999).

### Spawning is Influenced by Latitude and Location

The amount of solar power (irradiance) and the impacts of daily solar energy (insolation) are important to coral reproductive cycles as the energy corals receive through photosynthesis is a key part of their spawning behavior. Understanding the interplay of insolation and spawning behavior can help define spawning triggers in *Orbicella* spp. and identify the role of light in coral spawning. Spawning amount records indicate that, while Minimal spawning events occur as early as 4 and 5 DAFM, the primary spawning nights are synchronized within each species.

The time (HPS) of Peak spawning events did not vary with latitude, suggesting that spawning hour is not impacted by factors such as insolation. Observed variation in spawning time may be due to the local environment and annual or regional changes, supporting our hypothesis that the angle of the sun and total daily insolation is not the key driver of coral spawning hour. Our findings show that spawn month exhibits a significant gradient based on latitude (p<0.05), suggesting that the insolation may impact the overall reproductive cycle and large-scale developmental processes. Additionally, we found that Peak spawning date (DAFM) was significantly correlated with latitude in *O. annularis* and *O. faveolata*, with a slight gradient exhibited in *O. franksi*, suggesting that spawn date is impacted by changes in lunar light levels associated with latitudinal gradients. Light is crucial for circadian rhythms and maintenance of biological clocks, and the start of coral reproductive cycles may be directly tied to broad changes in light level that occur seasonally due to shifts in Earth’s angle to the sun (Rosenberg et al., 2017). Seasonal changes in energy availability may initiate reproductive cycles at different times between the Southern and Northern Caribbean locations, leading to different months of gamete release as observed in this study. Spawning hour is consistent over latitudinal gradients and varies within each geographic location, suggesting that it may be impacted more by environmental conditions at a local scale.

The lack of correlation between spawning time and insolation is further supported by our analysis of spawning hour in relation to colony depth as we found no significant change in spawning hour between colonies at the top and bottom of that species’ depth range. Our data capture the typical depth ranges of each species, with *O. annularis* ranging from approximately 3-20m, *O. faveolata* ranging from approximately 3-40m, and *O. franksi* approximately 12-20m. It should be noted that many observers did not record colony depth for each colony, and either noted an average depth for their collection or no depth at all. While this may limit the certainty of our findings, Levitan et al. (2011) have reported that depth does not appear significantly impact the spawning time of *O. annularis* and *O. franksi* in Panama. Their findings support our findings and further suggest that solar irradiance does not act as a direct trigger of spawning time in *Orbicella* species.

Increased understanding of spawning behavior and factors that lead to spawning synchrony is crucial to mitigate spawning desynchronization. Our study provides a valuable baseline of *Orbicella* spp. reproductive timing and is a resource for future work relating light and time of spawning. Recent studies indicate that spawning time is impacted by artificial light at night (ALAN) and reefs closer to large urban developments exhibit earlier spawning times (Ayalon et al., 2020; Davies et al., 2023). Regional temperature increases can generate similar spawning desynchronization or lead to reduced spawning in affected areas (Shlesinger & Loya, 2019; Monfared et al., 2023; Sheridan et al., 2025). The observed regional desynchronization at these sites could allow for more frequent cross-fertilization in those locations in species like *O. annularis* and *O. franksi* or limit intra-species fertilization success if differences in reproductive timing lower gamete density in the water column (Davies et al., 2023). Our findings indicate that observed variability in spawning date and time may be an entrained response resulting from both circadian rhythms and location-specific environmental factors. Our meta-analysis provides a baseline for future studies looking at perturbations to typical reproductive behavior in the *Orbicella* species complex.

## Notes

### Competing Interest Statement

The authors have declared no competing interest.

